# Differential diffusional properties in loose and tight docking prior to membrane fusion

**DOI:** 10.1101/2020.03.28.013482

**Authors:** Agata Witkowska, Susann Spindler, Reza Gholami Mahmoodabadi, Vahid Sandoghdar, Reinhard Jahn

**Author notes:** **Correspondence:** * (R.J.), (V.S.), and (A.W.).

## Abstract

Fusion of biological membranes, although mediated by divergent proteins, is believed to follow a common pathway. It proceeds through distinct steps including docking, merger of proximal leaflets (stalk formation), and formation of a fusion pore. However, the structure of these intermediates is difficult to study due to their short lifetime. Previously, we observed a loosely and tightly docked state preceding leaflet merger using arresting point mutations in SNARE proteins, but the nature of these states remained elusive. Here we used interferometric scattering (iSCAT) microscopy to monitor diffusion of single vesicles across the surface of giant unilamellar vesicles (GUVs). We observed that the diffusion coefficients of arrested vesicles decreased during progression through the intermediate states. Modeling allowed for predicting the number of tethering SNARE complexes upon loose docking and the size of the interacting membrane patches upon tight docking. These results shed new light on the nature of membrane-membrane interactions immediately before fusion.

## Introduction

SNARE (soluble N-ethylmaleimide-sensitive factor activating protein receptor) proteins mediate membrane fusion in the secretory pathway of eukaryotic cells and are widely used as model for studying fusion. They are small, mostly membrane-anchored proteins characterized by a conserved motif (SNARE-motif) that is usually located adjacent to the C-terminal membrane anchor (1). Upon contact between membranes destined to fuse, four complementary SNARE motifs assemble to form a trans-SNARE complex that cross-links the membranes. Assembly is initiated at the membrane-distal ends and then progresses towards the C-terminal membrane anchors (“zippering”), thus pulling the membranes tightly together (1–3). Zippering is highly exergonic and thus overcomes the energy barriers for membrane fusion (4–7). After membrane merger is completed, all SNAREs of the complex are aligned in parallel in the same membrane (cis-SNARE complex).

Despite major progress, the structural transitions along the fusion pathway are far from clear. Fusion commences with membrane contact and then leads via non-bilayer transition states to the final opening of an aqueous fusion pore. Experimental evidence complemented with both continuum and particle based models have suggested tentative pathways involving tight membrane contact brought about by SNARE zippering, associated with local protrusions of high curvature in some models (8–10). This state may then lead to lipid tail splaying between the fusing bilayers (11), resulting in a merger of the proximal monolayers yielding a structure that is also referred to as fusion stalk (see e.g. ref. 12). From the stalk intermediate, the system may progress to forming a hemifusion diaphragm (13) before the final opening of the fusion pore (8). For the transition from one intermediate to the next, energy barriers such as electrostatic repulsion or steric hydration forces need to be overcome (14–16).

Under physiological conditions, the lifetime of these intermediate states is very short. Indeed, only recently docking intermediates preceding hemifusion in SNARE-mediated fusion have been captured by cryo-electron microscopy (6, 17, 18). Deletion of a single amino acid in the SNARE synaptobrevin (Δ84 syb) results in vesicles that are arrested in a tightly docked (but not hemifused) state in which the membranes are separated by less than one nanometer. In contrast, substitution of two amino acids more distal from the transmembrane domain (I45A,M46A; referred to as AA syb) uncovered a loosely docked state where the membranes are separated by a larger gap, only connected by reversible SNARE interactions (17). Finally, extended hemifusion diaphragms were observed when the wild type (WT) syb protein was used (6). In this study, we have used *in vitro* reconstitution in order to characterize the nature of the loose and tight docking states. In particular, we have analyzed whether vesicles arrested in one of these states are capable of diffusing on the surface of the target membrane, and whether diffusion is affected by the nature of the docking intermediate. To this end, we have taken advantage of a previously established model system in which the SNARE-mediated docking and fusion of small vesicles with a single giant unilamellar vesicle (GUV) can be measured (19). In this work, we used the neuronal SNARE proteins synaptobrevin-2, syntaxin-1, and SNAP-25 as model, with synaptobrevin incorporated into small vesicles and an activated acceptor complex of syntaxin and SNAP-25 (20) inserted into the GUV membrane. This system is ideally suited for studying particle diffusion on membranes because the membrane area is large while diffusion is not hindered by any type of surface contact as, for instance, in supported planar bilayers (21). We employed interferometric scattering (iSCAT) microscopy to record three-dimensional (3D) trajectories of particle diffusion on the surface of GUVs. This technique has been shown to reach microsecond temporal resolution and nm precision in detecting and tracking individual nanoparticles via the interference of their scattered light with a reference beam (22, 23, and see Methods). Using iSCAT, we could investigate the diffusion of unlabeled vesicles arrested at a loosely or tightly docked intermediate state using synaptobrevin point mutants as previously described (17). Modeling of the diffusion data allowed for approximating the number of SNARE complexes involved in vesicle docking (loose docking) as well as the size of the interaction interface (tight docking).

## Results

To study loosely and tightly docked intermediates, a GUV containing SNARE acceptor complexes was immobilized with a micropipette (see Figure 1a) and brought in close proximity to the cover glass. Next, we added large unilamellar vesicles (LUVs, 80–100nm in diameter) containing synaptobrevin (syb) and used iSCAT microscopy to track LUVs on the lower hemisphere of individual GUVs from a single focus position (22). The LUVs readily bound to the GUV and either fused with it (as for WT syb, ref. 19) or they remained docked on its surface and diffused. Detection of the LUV scattering signal was greatly improved with a background-correction procedure (compare upper and lower image in Figure 1b) described in more detail in Methods. The iSCAT contrast of a nano-object is directly proportional to its polarizability, which is in turn linked to the particle size (23). The size of the LUVs was estimated from the observed iSCAT contrast to be about 90 nm (see Figure 1c). Thus, an LUV appears as the iSCAT point spread function (PSF) of microscope, which consists of few lobes encoding the 3D position of the scatterer (24). This PSF could be fitted with a physical model to extract the third dimension (25), however here we fitted the main lobe for the lateral localization and used the geometry of the GUVs to extract the axial position of the vesicles (Figure 1b).

**Figure 1.**
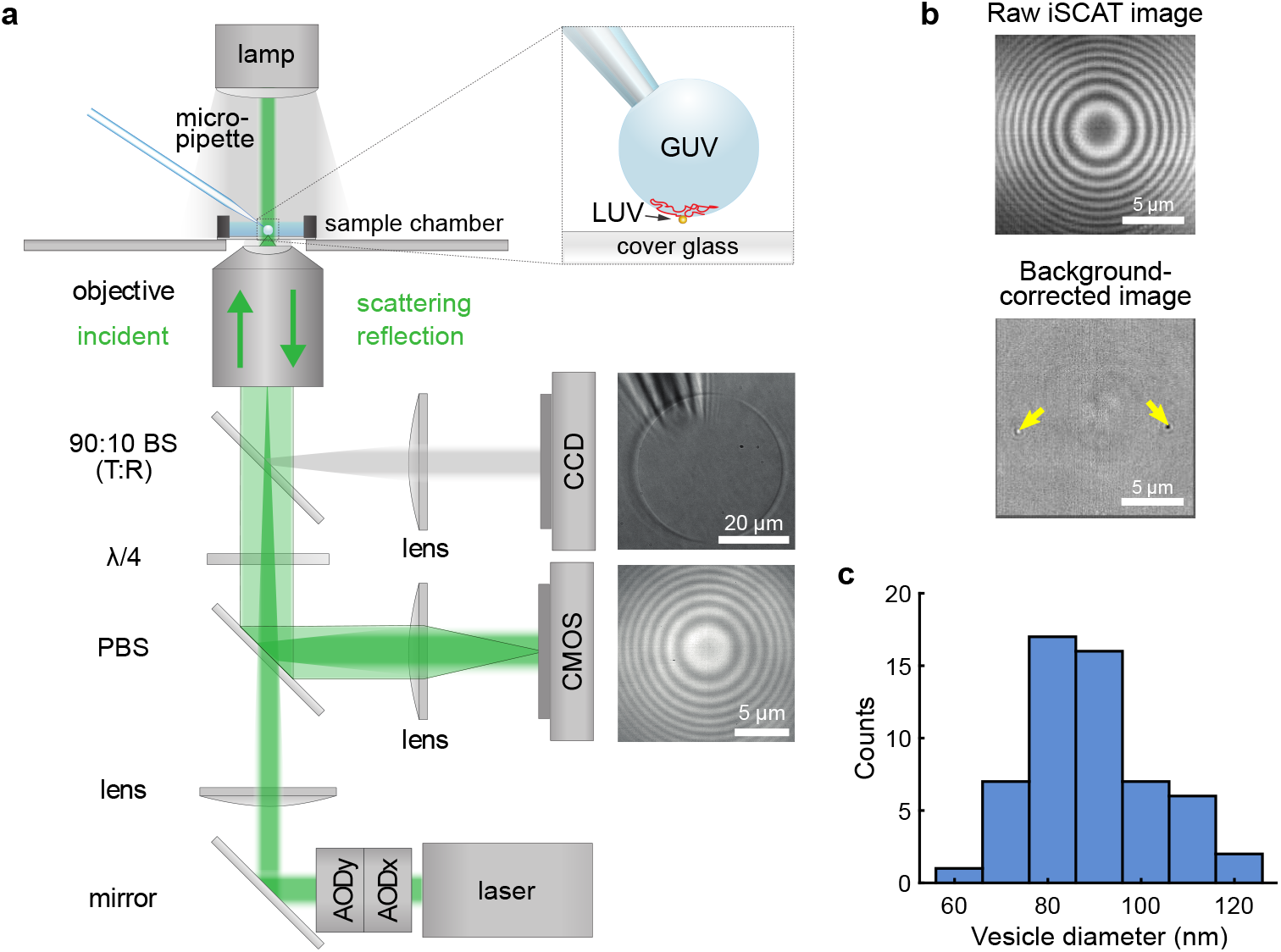
iSCAT detection of GUVs and liposomes. (a) Schematic illustration of the experimental optical setup allowing the observation of GUVs in bright field (on the CCD camera) and iSCAT (on the CMOS camera) as well as micropipette manipulation of GUVs (see inset). BS—beam splitter, λ/4—quarter wave plate, PBS—polarizing beam splitter, AOD—acousto-optical deflector. (b) iSCAT GUV image before (upper image) and after (lower image) background correction procedure. In the background-corrected image, two docked LUVs become visible (marked with yellow arrows) with the PSF shape of the wide-field iSCAT microscope encoding the particular height of the particles. (c) Histogram of average LUV size determined based on their contrast in iSCAT.

Lack of photobleaching in iSCAT allowed for monitoring the 3D motion of docked vesicles over prolonged periods (as shown exemplarily in Figure 2a) while maintaining nanometer spatial precision. Docked vesicles closely follow the topography of the GUV surface, thus displaying a characteristic periodic ring-shaped contrast pattern (see Figure 2b), similar to the Newton rings encountered in Figure 1b (also see inset in Figure 2b). We then compared surface diffusion of docked vesicles containing either WT syb or the docking mutants Δ84 syb and AA syb (described above). As expected, all three vesicle populations dock to GUVs that contain acceptor SNARE complexes. However, when we determined their diffusion coefficients from the measured 3D trajectories (Figure 2a), significant differences were observed between the three populations (Figure 2c). Vesicles carrying the AA syb mutants showed the fastest diffusion, followed by those containing the Δ84 mutant whereas vesicles with WT protein were the slowest. These data show that the different states uncovered previously by electron microscopy (17) indeed represent physical differences in the nature of membrane attachment between the two vesicles.

**Figure 2.**
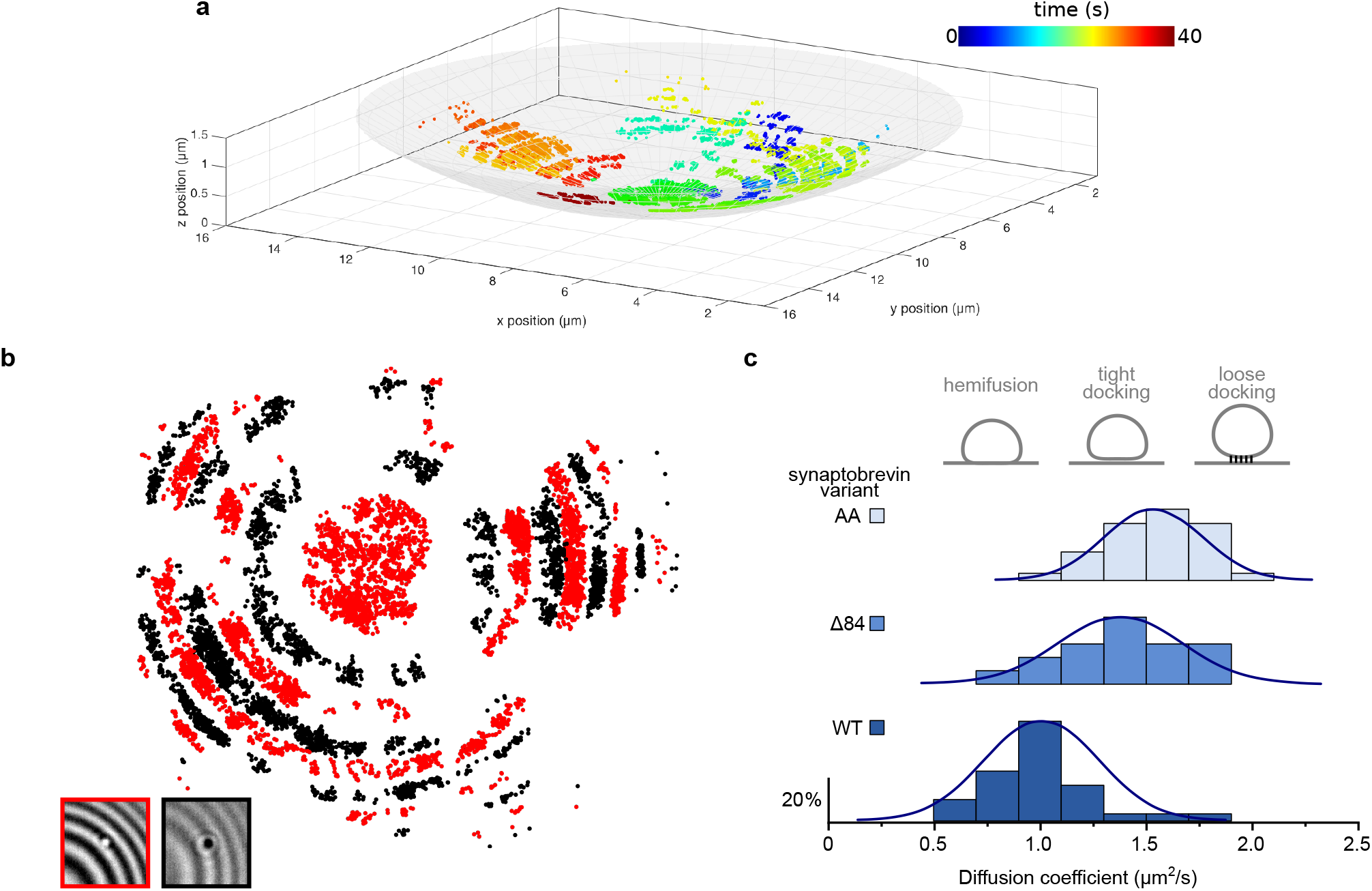
Tracking of docked LUVs on a GUV surface. (a) 3D reconstructed trajectory of a single liposome docked on the GUV surface tracked for 40 s. The color bar shows the time evolution. The sphere represents a GUV surface with a radius of 18 μm (b) *xy* projection of the docked liposome’s trajectory from panel a. The point color depends on the LUV iSCAT signal intensity reflecting fluctuations in the LUV z-position relative to the GUV membrane: Regions marked in black and red correspond to the regions of dark and bright iSCAT particle contrasts, respectively (see lower insets). The gaps in the trajectory between regions with positive and negative contrast result from the fact that the vesicle is barely visible when undergoing a contrast switch. (c) Histograms presenting diffusion coefficients of LUVs docked on the GUV surface depending on the synaptobrevin variant (AA, Δ84, or WT). Bin width 0.2μm^2^/s. Blue lines represent a Gaussian fit. N_AA_=32, N_Δ84_=17, N_WT_=32. Illustrations above the graph depict possible docking states of the vesicles.

As discussed above, the loosely docked state is probably governed exclusively by trans-interactions between individual SNAREs whereas in the tightly docked state the membranes are firmly attached, forming a disk that is probably rigid (6, 17). Using the experimentally obtained diffusion data, we have therefore carried out simulations in order to evaluate whether the data are compatible with these models. First, we modeled the diffusion of vesicles connected by multiple (1–20) SNARE complexes (Figure 3a upper panel). To gain insight into the diffusion of a liposome docked on the GUV surface while being tethered to different numbers of SNARE complexes, we simulated random walks of individual SNARE complexes within the constraining circular area to reflect the maximal docking interface area (Figure 3b, for details see Methods). Vesicle position was repeatedly determined after each displacement step as center of mass of all simulated proteins (Figure 3b) to form a 5 s-long trajectory (see Figure 3c). Next, mean squared displacement (MSD) curves were generated from the trajectories (Figure 3d), and diffusion coefficients for varying number of tethering SNARE complexes were calculated (Figure 3e). The outcome of simulations confirms that the diffusion coefficient of a vesicle decreases with increasing number of tethering SNARE complexes (Figure 3c and e). Diffusion speeds obtained in simulations and experiments let us conclude that loosely docked vesicles (mainly AA syb and Δ84 syb) observed with iSCAT are attached to the GUV with maximally 3–4 assembled SNARE complexes. Docked vesicles moving at the maximal speed, on the other hand, are found to be attached via 2 SNARE complexes. Vesicles attached with 1 SNARE complex were probably rejected from our analyses due to large fluctuations in position perpendicular to the GUV surface and lack of a ring-shaped amplitude pattern as seen in Figure 2b.

**Figure 3.**
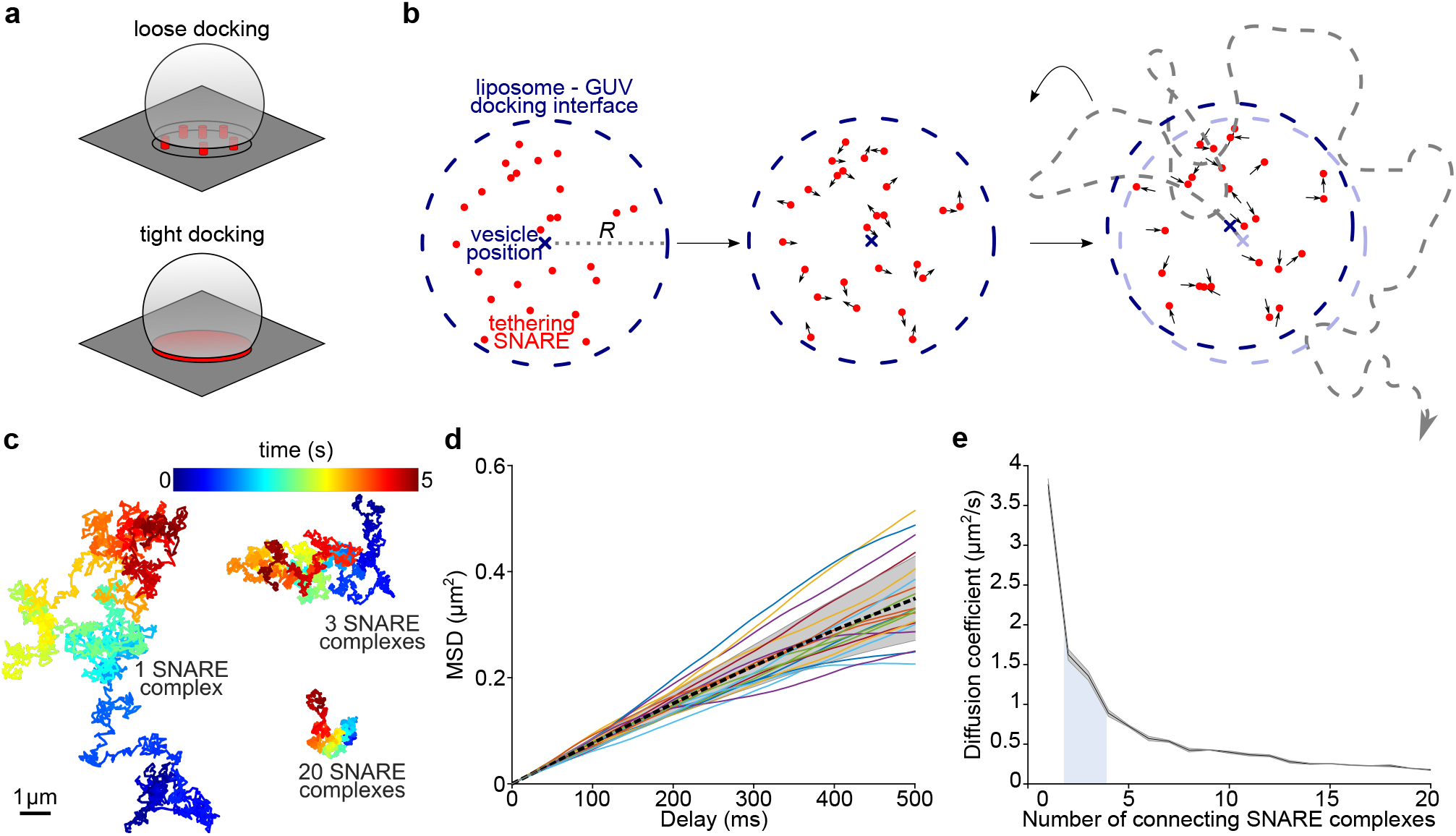
Simulation of diffusion of loosely docked vesicles tethered by multiple SNARE complexes. (a) Schematic illustration of a vesicle tethered by multiple SNARE complexes to GUV membrane (loose docking) and vesicle that is tightly docked on a GUV surface with interacting docking interfaces marked in red. (b) Schematic illustration of the random walk diffusion simulation of SNARE proteins (red points) with liposome position determination (cross) and with a diffusional constraint *R* for SNARE proteins representing maximal contact area between LUV and GUV (dashed blue line). Liposome trajectory after multiple displacement steps is schematically shown with a grey dashed line. (c) Representative trajectories obtained in simulations with 1,3, or 20 tethering SNARE complexes. (d) Mean square displacement (MSD) obtained from 20 simulation runs with 20 tethering SNARE complexes. The mean is represented by the dashed black line and the standard deviation of the mean is shown by the shaded area. (e) Dependence of LUV’s diffusion coefficient on the number of tethering complexes. The mean is represented by the black line and the standard deviation of the mean is shown by the shaded area around it. Each diffusion coefficient was determined in 20 independent simulation runs. The blue area below the graph shows the range of experimentally measured diffusion values for synaptobrevin Δ84 and AA.

Next, we modelled the diffusion of vesicles that were attached to the GUV through tight docking interfaces by assuming a disk-like docking interface that diffuses as a whole (see Methods). Here, we have used an analytical approximation of the translational diffusion in the membrane (26, 27) to calculate the theoretical diffusion coefficient of a tightly docked vesicle with a given docking interface radius (Figure 4a). The resulting diffusion coefficients (Figure 4c) decrease with increasing interface size at a slower pace than in the case of a loose docking model (Figure 3e). Remarkably, diffusion coefficient values of the slowest vesicles containing syb mutants (i.e., potentially not hemifused, Figure 2c) correspond to the modelled interface radius of around 50 nm, which, in turn, correlates with maximal feasible LUV deformations based on electron microscopy studies (6, 17).

**Figure 4.**
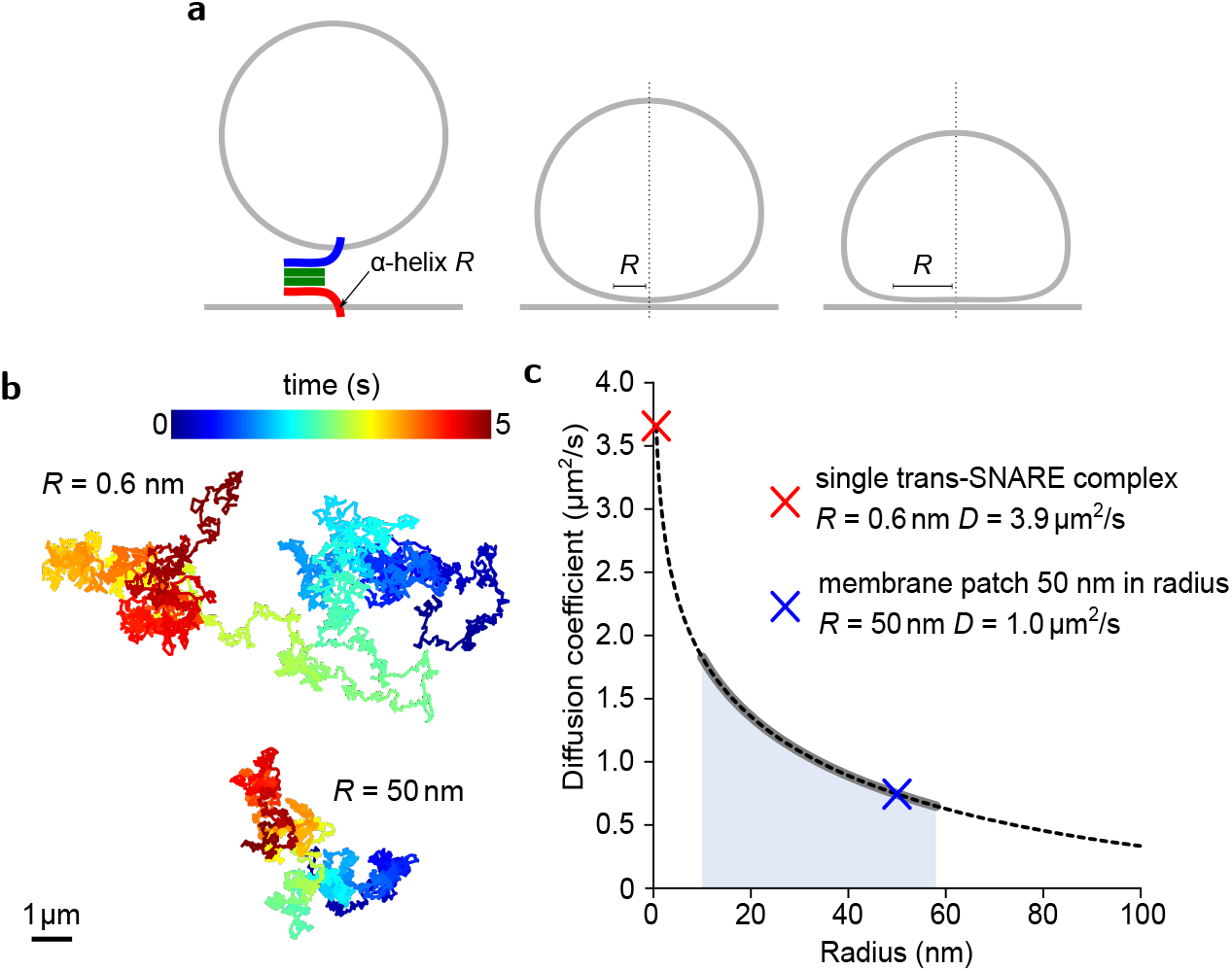
Modelling the diffusion of tightly docked vesicles. (a) Schematic illustration showing different docking stages that can be captured by the model: point attachment by a single tethering SNARE complex, intermediate, and extended docking interfaces with varying radii *R* of the membrane interaction surface. (b) Example trajectories of vesicles over 5 s, moving with diffusion speeds obtained from the model (see panel c) corresponding to point attachment (R = 0.6nm) and extended docking interface (R = 50 nm). (c) Analytical solution of the tight docking model (dashed line) with parameters (Table 2) corresponding to the experimental conditions in this work. Thickened grey line and blue area show the range of experimentally measured diffusion values for synapto-brevin Δ84 and AA. Diffusion coefficient values (D) of the trajectories shown in panel b are marked with crosses.

## Discussion

Here we utilized *in vitro* reconstitution using previously described SNARE mutants (6, 17) in order to characterize recently discovered early membrane fusion intermediates (6, 17). Employing iSCAT microscopy in this work allowed for the first time for extended 3D tracking of vesicles’ dynamics on large, free-standing membranes with very low curvature as typical for cell membranes, thus avoiding disturbance by surface contact or edge effects that may occur in supported or pore-spanning membranes (28). With this approach we were able to observe that docked vesicle’s arrest at subsequent membrane fusion intermediate states (loose docking, tight docking, and hemifusion) is reflected in decreasing diffusion coefficients.

The loosely docked state, stabilized here by point mutations in the SNARE synaptobrevin, may serve as model for partially assembled SNARE complexes in which progression of zippering is thought to be arrested by regulatory proteins (1, 29). In our experimental system, we find that such loosely docked vesicles on average diffuse faster than other docked liposomes which indicates less interactions between docked membranes. Moreover, the number of connecting SNARE complexes predicted by our model (up to 3–4) is in good agreement with current, albeit rather indirect, estimates for the number of SNARE complexes required for fusion (30–32).

Tightly docked membranes have only recently been observed in fusion reactions mediated by SNARE proteins (6), viral (33) and mitochondrial fusion proteins (34). Additionally, this state appears to be also independent of the continued presence of the tethering proteins, since docking becomes irreversible upon SNARE complex disassembly (17). Moreover, development of tight membrane-membrane contact patch was also inferred from protein-free coarse-grain simulations (35), providing further evidence that it constitutes a true fusion intermediate. Taken together, it seems that tight docking is independent of the nature of the fusion protein, and is thus an intrinsic feature of the bilayer. In this work, we detected a high proportion of vesicles with estimated relatively large tight attachment interfaces, exceeding 10 nm in radius. Together with previous electron microscopy observations of large, flattened, interacting membrane patches (6, 33, 34), it suggests that once formed, expansion of such an adhesion interface can progress up to the point when further membrane deformation is not possible or when there is steric hindrance coming from protein tethers. It remains to be established how the repulsive forces of the negatively charged membranes are overcome at close distance, and also whether, and to which extent, diffusion of membrane constituents is hindered in the disk-like contact zone. Do tightly-docked intermediates represent fusion intermediates in living cells? Extended disk-shaped contacts were frequently observed in classical electron microscopy studies of secretory vesicles (36), but they were later attributed to fixation artefacts (37). However, more recently docked vesicles with sub-nanometer distance from the plasma membrane were seen in electron microscopy images of synaptic vesicles in neurons obtained with high-pressure freezing (38). Understanding the physical parameters governing this state such as adhesive and repulsive forces at the contact zone, the lateral mobility of molecules within and between the membranes, and the energy barriers leading from this state to nonbilayer-intermediates such as fusion stalks will be instrumental for unraveling the mechanism of membrane fusion. The methodology applied here may be useful for analyzing the intermediate states of other fusion proteins such as those mediating mitochondrial or viral fusion.

## Methods

### Protein expression and purification

*Rattus norvegicus-derived* SNARE proteins (syntaxin-1A (183–288) (39), SNAP-25 (cysteine free) (40), synaptobrevin-2 (wildtype (WT) (41), Δ84 (42), and I45A,M46A (AA) (17, 43)), 1–96 (44), and synaptobrevin-2 fragment (49–96) (20) were expressed and purified as described before (17, 45). In short, proteins were expressed in *Escherichia coli* strain BL21 (DE3) and purified via nickel-nitrilotriacetic acid affinity chromatography (Qiagen) followed by ion exchange chromatography on an Äkta system (GE Healthcare). The preassembled so called ΔN complex (20) was used to mimic plasma membrane SNARE acceptor complex. It consists of syntaxin, SNAP-25, and synaptobrevin fragment 49–96. The complex was obtained by mixing the monomers overnight at 4°C, followed by purification of the complex using ion exchange chromatography (MonoQ column) in a buffer containing CHAPS as described before (20, 45).

### Liposome preparation

Lipids—brain-derived PC, PE, and PS along with cholesterol (ovine wool)—were purchased from Avanti Polar Lipids. For all liposome mixtures lipids PC, PE, PS, and cholesterol were mixed in a ratio of 5:2:2:1, respectively.

Large unilamellar vesicles (LUVs, diameter ~100 nm) were prepared with a reverse phase evaporation method as previously described (6, 19). Proteoliposomes were formed using a direct reconstitution method as previously described (6), with synaptobrevin WT, Δ84, or AA reconstituted in a protein/lipid ratio of 1:500.

Small proteoliposomes (diameter ~40nm) were prepared as described before (19) by co-micellization followed by size exclusion chromatography. During preparation ΔN complex was added to yield a protein/lipid ratio of 1:1000.

### GUV preparation

Giant unilamellar vesicles were prepared from small proteoliposomes containing ΔN complex by electroformation using in-house-built electroformation chambers containing platinum wires (46) using a protocol described previously (45).

### iSCAT measurements

Liposomes were tracked using iSCAT microscopy (47). The basic principle is schematically shown in Figure 1a (a more detailed description of the setup can be found in ref. 28). Briefly, the incoming light *E*_inc_ of a 532 nm laser (Verdi 2G, Coherent) is scattered from the GUV and the liposomes (*E*_scat_) and partially reflected at the glass—sample interface (*E*_ref_). *E*_scat_ and *E*_ref_ are collected via an oil-immersion 60× objective (Olympus ApoN, NA = 1.49) and imaged onto a CMOS camera (MV-D1024E-CL, Photonfocus AG). The intensity at the camera is given by

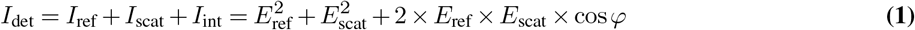

and is thus dependent on the phase *φ* between the scatterer and the reflected light. Changes in the distance between the scatterer and the glass—sample interface result in a periodic modulation of the scatterer’s contrast. Scattering at the GUV surface therefore creates a ring pattern, as shown in Figure 1b. In the same fashion, the point spread function of a liposome changes its amplitude when it travels along the GUV surface and thus appears either bright or dark on the GUV background.

To hold the GUVs in place above the field of view, we used micropipette aspiration (48) using glass pipettes with ~5 μm opening connected to a height-adjustable water tank and a micromanipulator (Sensapex). Using a 90:10 (T:R) beam splitter, a part of the collected light was sent onto a CCD camera (sensicam qe, PCO) for monitoring the GUV aspiration process in bright field.

Videos of liposomes bound to GUVs were recorded typically for 5 s at 1 kHz frame rate. For liposome tracking, we performed background correction based on image registration. The iSCAT contrast of the liposomes typically amounts to 2–3%. In the background corrected images, we fit a 2D Gaussian function to each point spread function for determining the 2D projected position. The particle height *z_i_* in each frame is calculated based on the GUV center (*x*_GUV_,*y*_GUV_), the GUV radius *r*_GUV_, which we extract from the bright field image, and the 2D particle localization (*x_i_, y_i_*):

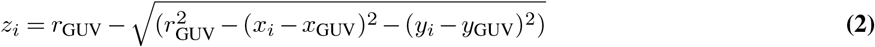

We determined the center of the GUV using two consecutive routines adopted from (49). The diffusion coefficient was calculated for each trajectory from the first two points of the mean squared displacement (MSD(*τ*)) with time lag *τ* of Δ*t* and 2Δ*t* (Δ*t* = 1 ms):

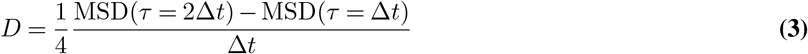

### Modelling of loose docking

Liposomes loosely docked on the GUV surface were assumed to be tethered by different numbers of SNARE complexes. Their diffusion was simulated using random walk (50) of individual SNARE complexes in a 2-dimentional membrane (similar to ref. 51). The constraining circular area (50 nm in radius) was introduced to reflect the maximal surface of the liposome-GUV docking interface with size resembling maximal LUV membrane deformation during docking as seen in (6, 17, 32). Other radii (5–100 nm) of constraint areas including adaptive one (constraint radius increasing with increasing number of tethering SNARE complexes) were also tested, however did not yield substantially different results (data not shown). The effect of solvent friction on vesicle’s diffusion in this case was assumed to be negligible. It also seems unlikely that membrane deformation generated here by diffusing SNARE complexes would result in major decrease of the diffusion coefficient. Therefore, we did not introduce such energy potential in our model. Initially, various number of tethering SNARE complexes are randomly distributed within the constraining area. Each SNARE complex displacement *dx* was determined in every time step *dt* (usually 100 μs):

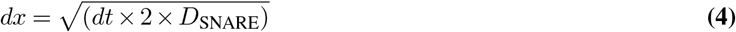

with *D_SNARE_* being the diffusion coefficient of the assembled SNARE complex of 3.8 μm^2^/s as determined in (52). The absolute position of a vesicle after each displacement step was then calculated as center of mass of all simulated proteins. Each 5 s-long simulation was performed 20 × to determine average MSD and diffusion coefficient of the vesicle. Further, we have investigated how clustering of SNARE complexes influences diffusion of loosely docked vesicles, but we did not observe any large effects (not shown). The source code used for simulations in Octave (53) and MATLAB (MathWorks) is available at Zenodo (54).

### Modelling of tight docking

The diffusion speed of tightly docked vesicles was modelled with the use of an analytical expression (26) for the model developed earlier by Hughes, Pailthorpe, and White (27) to predict diffusion of a membrane embedded objects (proteins) based on the hydrodynamic radius of the membrane embedded part. The version of the approximation used in this study was presented in ref. 55:

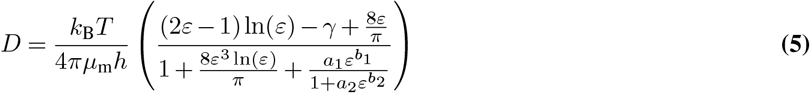

with the parameter *ε* being dependent on the inclusion radius *R*:

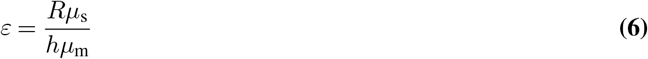

Explanation of used symbols can be found in Table 1. Values used in the model are given in Table 2. Simulated example trajectories of tightly docked vesicles were generated as random walk (Equation 4) of objects of certain radius characterized by the diffusion coefficient obtained from Equation 5.

**Table 1.** Symbols used in Equation 5 and 6

| Symbol | Explanation |
| --- | --- |
| $D$ | diffusion coefficient |
| $k_B$ | Boltzmann's constant |
| $T$ | temperature (in K) |
| $\mu_m$ | membrane viscosity (in Pa × s) |
| $h$ | bilayer thickness |
| $\gamma$ | Euler's constant |
| $\varepsilon$ | parameter derived from Equation 6 |
| $a_1, b_1, a_2, b_2$ | equation scaling coefficients (56) |
| $R$ | membrane inclusion radius |
| $\mu_s$ | viscosity of surrounding medium (in Pa × s) |

**Table 2.** Model parameter values used in Equation 5 and 6

| Parameter | Value | Explanation |
| --- | --- | --- |
| $k_B$ | $1.38064852 \times 10^{-23} \frac{\text{m}^2 \times \text{kg}}{\text{s}^2 \times \text{K}}$ | physical constant |
| $T$ | 293 K | $\approx 20^\circ \text{C}$ |
| $\mu_m$ | 115 mPa × s | calculated from fastest measured diffusion coefficient $\approx 3.9 \frac{\mu\text{m}^2}{\text{s}}$ determined by FCS in ref. 52 |
| $h$ | 4.6 nm | as in ref. 51 |
| $\gamma$ | 0.5772 | physical constant |
| $a_1$ | 0.433274 | 56 |
| $b_1$ | 2.74819 | 56 |
| $a_2$ | 0.670045 | 56 |
| $b_2$ | 0.614465 | 56 |
| $R$ | 0.6 nm | for single SNARE complex transmembrane domain (57) |
| $\mu_s$ | 0.96 mPa × s | 55 |

## AUTHOR CONTRIBUTIONS

A.W., R.J., and V.S. designed the study. A.W. prepared all samples, performed confocal imaging, wrote code and performed numerical simulations, analyzed and evaluated all data. S.S. prepared GUVs, performed and evaluated iSCAT experiments. R.G.M. contributed new analytic tools for iSCAT, evaluated iSCAT experiments. A.W., S.S., R.J., and V.S. wrote the manuscript.

## ACKNOWLEDGEMENTS

We would like to thank Martin Kaller for support with iSCAT experiments. This work was supported by funds from the Max Planck Society (to R.J. and V.S.), US National Institutes of Health grant No. 2 P01 GM072694 (to R.J.), and an Alexander von Humboldt Professorship (to V.S.). For this preprint we adapted template from the group of Dr. Ricardo Henriques.

